# EndoGenius: Enabling comprehensive identification and quantitation of endogenous peptides

**DOI:** 10.1101/2025.06.12.659347

**Authors:** Lauren Fields, Tina C. Dang, Mitchell Gray, Satirtha S. Protya, Lingjun Li

**Affiliations:** Department of Chemistry, University of Wisconsin-Madison, 1101 University Avenue, Madison, WI 53706; School of Pharmacy, University of Wisconsin-Madison, 777 Highland Avenue, Madison, WI 53705

## Abstract

**Summary:** The investigation of endogenous peptides, specifically with respect to neuropeptides, from mass spectrometry data is rife with bioinformatics bottlenecks, stemming from the low *in vivo* abundance of these analytes, increased susceptibility to degradation, and an immense search space of possible peptides. To address this, we present EndoGenius in its expanded form, strategically designed to optimize the searching for these endogenous peptides complemented with a pipeline designed for tasks including quantitation, spectral library building, motif extraction, and usage with data-independent acquisition workflows.

**Availability and Implementation:** EndoGenius is released as an open-source software package under an MIT License. The EndoGenius package with a user interface can be installed from https://www.lilabs.org/resources. The source code for EndoGenius can be accessed at https://github.com/lingjunli-research/EndoGenius-v2.0.

## Introduction

Neuropeptides represent a class of endogenous signaling molecules with the unique ability to provide a dynamic fingerprint of *in vivo* neuronal functions. Despite this potential, neuropeptides are often overlooked, in part due to the complexities that arise through signaling cascades. Thus, the crustacean model has been adopted as a premier test bed for the investigation of neuropeptides,(Christie, et al., 2010; Marder and Bucher, 2007) through which the physiological response to feeding,(Turrigiano and Selverston, 1990) hypoxia,(Chung and Webster, 2005) and copper toxicity,(Sauer and Li, 2021) and others have been evaluated. Even in this simpler model system, however, the bioinformatics challenges that accompany analysis of neuropeptides were, until recently, unaddressed. As neuropeptides are present in a wide range of sizes, from three residues to well over one hundred residues in length, and the endogenous cleavage patterns of these molecules are largely unknown, these biological conditions are suboptimal for the typical bottom-up proteomics workflow, which often uses enzymatic digestion producing a smaller search space of ions with higher ionization efficiency. Without this advantage, neuropeptides must be evaluated in a digest free manner.(De la Toba, et al., 2022) This is coupled with their low *in vivo* abundance and high propensity to degradation, further adding challenges both experimentally and computationally.(Maes, et al., 2014)

We recently announced our new software suite, EndoGenius, which is designed to address all of the unique bioinformatics challenges that accompany analysis of neuropeptides, and endogenous peptides in general, with mass spectrometry. Through development of a novel scoring algorithm, in combination with leveraging conserved motif information, we were able to provide a software program able to outperform more traditional packages designed for proteomics. Herein, we describe an expansive addition to the initially reported EndoGenius framework,(Fields, et al., 2024) fit to accommodate all experimental needs, including processing of DDA and DIA spectral datasets, quantitation in label-free and isobarically-labeled formats, and additional tools such as MotifQuest.(Dang, et al., 2024)

## Methods

EndoGenius was written in Python and comes equipped with a Windows-compatible user interface, written with Tkinter. The complete program comes with the database searching algorithm, as well as supplemental tools for support of MotifQuest integration, quantitation report generation, reporter ion extraction from isobarically-labelled data, and spectral library building. The latest version of EndoGenius is open source and available free of charge at GitHub, https://github.com/lingjunli-research/EndoGenius-v2.0, with an MIT License. This page also includes an in-depth user manual, ensuring ease of integration with any experimental design.

## Results

EndoGenius is designed to be a one-stop-shop for neuropeptide identifications in both DDA and DIA modes. In DDA mode, raw data can be uploaded in.MS2 and.mzML file formats using RawConverter (He, et al., 2015) and MSConvert (Chambers, et al., 2012) to perform file conversion, respectively. A complete depiction of the EndoGenius algorithm is depicted in **Figure 1**. From import, users provide a.FASTA-formatted peptide database of undigested peptides in a variety of lengths. A shuffle decoy entry is generated for each target entry. Upon calculation of corresponding metrics, including percent sequence coverage of fragments observed with respect to peptide length, and the spectral similarity to theoretical, conveyed by a hyperscore, an additional, novel motif score is applied. This motif score is obtained through reference of a motif database, reflecting regions of highly conserved peptide sequences, and aids in the parsing of peptides with high levels of sequence similarity. Initially, it was required that this motif database, a requirement for EndoGenius operation, was manually curated and provided for the execution of the program. However, we recently developed MotifQuest, which leverages its own statistical calculations coupled with multiple sequence alignment using ClustalOmega (Sievers and Higgins, 2018) to perform this task simply from the input of a single.FASTA file.(Dang, et al., 2024) In this release, MotifQuest is now fully integrated into EndoGenius, providing a seamless transition from pre-processing through data analysis (**Supplemental Figure 1**). This also expands to utility of EndoGenius beyond just neuropeptides to other endogenous peptides, including antimicrobial peptides.(Dang, et al., 2024) Following these filters, a scoring metric is applied from which the user can either calculate the false-discovery rate (FDR) associated with a particular score, filtering to the desired FDR threshold, or implement the EndoGenius score. For small datasets, in which less than one hundred identifications are expected, as is true with crustacean neuropeptides, it is recommended to apply an external scoring function, rather than FDR, as FDR can lack reproducibility in these sparse datasets. For these reasons, we integrate an EndoGenius score, which has been shown to have a linear relationship to FDR previously but does not succumb to the reproducibility issues present with FDR.(Fields, et al., 2024) With each EndoGenius run, a plot of FDR vs. EndoGenius score is provided, so that this relationship can be continually monitored.

**Figure 1:**
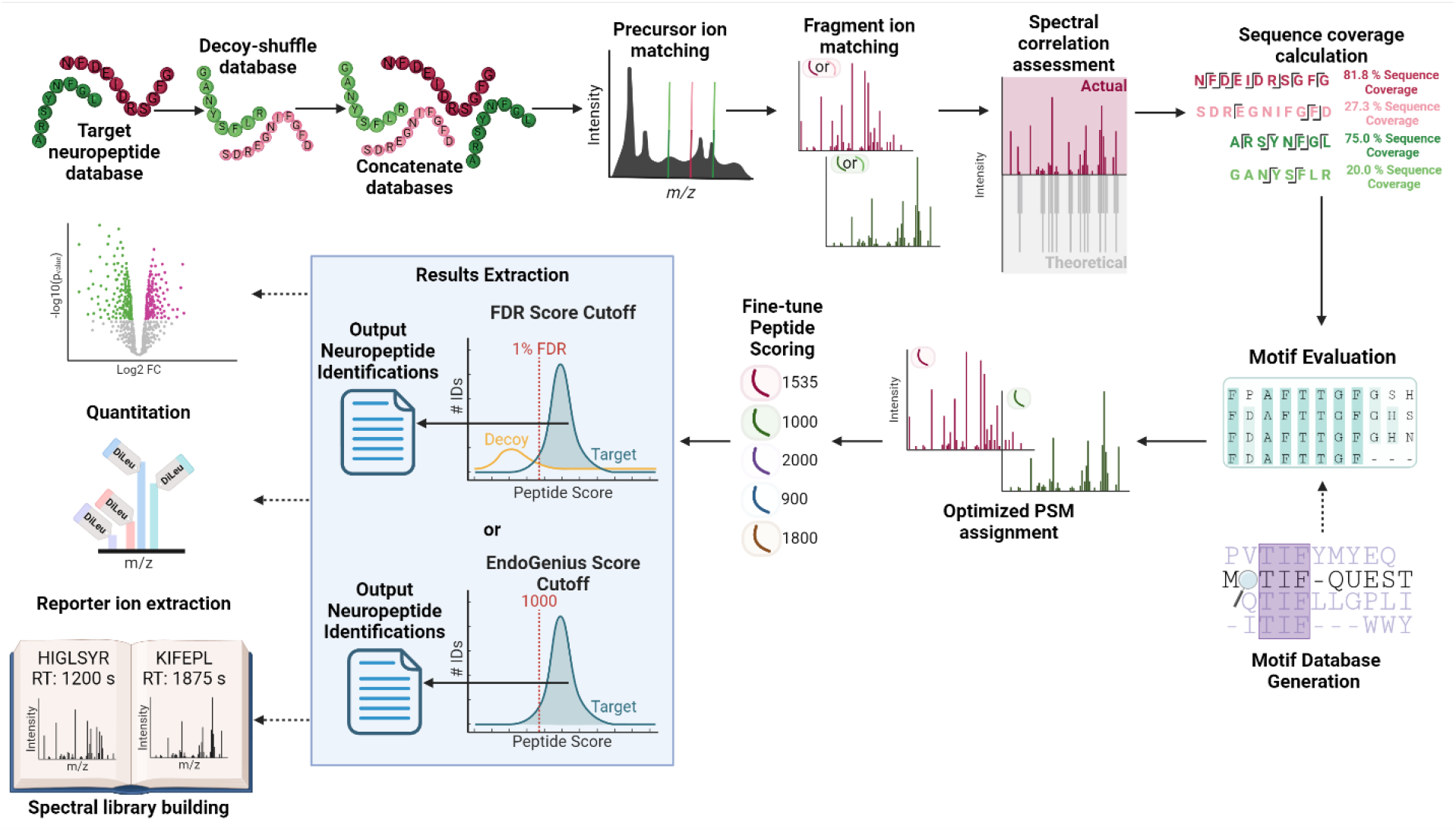
Improved workflow for EndoGenius. Option steps are denoted with a dashed arrow. First, spectra undergoes traditional database searching steps including precursor and fragment ion matching, spectral correlation, and sequence coverage calculation. Then, putative peptide-spectrum matches (PSMs) are evaluated for motifs, which can be achieved using the optional inclusion of MotifQuest software.(Dang, et al., 2024) PSMs are then finalized and scored. Following this, final results can be exported either using a false-discovery rate (FDR) or EndoGenius score as the quality threshold. Upon final identifications, a variety of supplemental operations can be conducted, including spectral library building, reporter ion extraction from isobarically-labeled sampled, and quantitation.

Following traditional database searching, quantitation is desirable. We sought to capitalize on the existing tools in our community, rather than regenerate something that already exists. To achieve this, we built an interface designed for utility in quantitation of label-free and isobarically-labeled peptides. With respect to label-free samples, the output reports from EndoGenius can be uploaded to the interface with annotation of sample names. All output reports are then merged according to peptide ID, providing the user with a report of all peptides IDed along with the precursor intensity, a common representation of peptide abundance used for quantitation. This report is formatted specifically for integration into the Perseus environment,(Tyanova, et al., 2016-06-27) wherein a myriad of statistical tools are available (**Supplemental Figure 2A**).

Additionally, it was desirable to provide support for isobarically-labeled tags, which afford the power of multiplexing, or analyzing many samples, sometimes twelve or more, in a single run. These samples are labeled with an isobaric reagent, and peptides, upon fragmentation, present a reporter ion in the low mass range of the MS/MS spectra.(Chen, et al., 2021; Paulo, 2022) These reporter ions are unique to each sample and can be used to relatively quantify between samples. To capitalize on this technology, we first integrated the inclusion of DiLeu as a variable post-translational modification (PTM) in the standard EndoGenius database search. Further, upon identification of peptides containing the DiLeu-modified N-termini, we provide an application in which the resulting report can be referenced to search for reporter ions for each peptide. These reporter ions are then able to be extracted and merged to provide a comprehensive intensity report for each peptide, fit for integration with Perseus, as before (**Supplemental Figure 2B**).

As neuropeptides are inherently of low abundance, use of DIA mass spectrometry methods has been shown to be advantageous in minimizing bias against these biomolecules of low abundance.(DeLaney and Li, 2019; Fields, et al., 2024; Phetsanthad, et al., 2023) To process complicated DIA spectra, in which often many precursor ions co-fragment, spectral libraries are often referenced to glean peptide identifications through comparison of library and DIA spectra. As spectral libraries are often generated from DDA spectra, we determined it would be ideal to equip EndoGenius to build spectral libraries from the optimized methods used to craft our results. To do this, we developed a built-in tool to EndoGenius that is able to merge an unlimited number of EndoGenius output files, and then generate a massive library of all identifications. As a spectral library typically should be restricted to just one entry from a particular peptide, we remove duplicate entries, retaining the peptide with the highest hyperscore, which reflects spectral similarity (**Supplemental Figure 3**). The output spectral library is then formatted in a.TSV file fit for integration with DIA-NN. DIA-NN is a neural-network based platform for processing DIA spectra, and is regarded for its efficiency and premier results.(Demichev, et al., 2020) The complete EndoGenius package is designed to be fully accessible for users of all experience levels, including a comprehensive GUI for the standard database searching (Supplemental Figure 4), which houses the utilities to fully customize all search parameters. As a pop-out from the main GUI, we support the integration of many post-translational modifications (PTMs), including 12-plex DiLeu (**Supplemental Figure 5**). From the tools menu, we then include interactive GUIs for use in performing quantitation (**Supplemental Figure 6A**), reporter ion extraction from isobarically-labeled samples (**Supplemental Figure 6B**), construction of spectral libraries from EndoGenius results (**Supplemental Figure 6C**), and operation of MotifQuest (**Supplemental Figure 6D**).

## Conclusions

Herein we present an expanded software suite designed specifically for the analysis of endogenous peptides, with use cases established in the areas of neuropeptidomics and antimicrobial peptidomics. This open-source software is designed specifically for the ability to seamlessly draw meaningful conclusions from peptidomics datasets and an optimized manner, with support for quantitation. Additionally, recognizing that DIA is perhaps a strategic way to obtain data from low abundance peptides, we incorporate the capability to build spectral libraries and then integrate DIA-NN for the interpretation of DIA datasets. Altogether, we present a unique platform tuned for complete peptidomic characterization.

## Acknowledgements

This work was supported in part by National Science Foundation (CHE-2108223) and National Institutes of Health (NIH) through grant R01DK071801. The Orbitrap instruments were purchased through the support of an NIH shared instrument grant (S10RR029531) and Office of the Vice Chancellor for Research and Graduate Education at the University of Wisconsin-Madison. T.C.D. was supported in part by National Institute of General Medical Services of the National Institute of Health under Award Number T32GM141013 (Molecular and Cellular Pharmacology Training Program) and the SciMed Graduate Research Scholars Fellowship through the University of Wisconsin-Madison. Support for this fellowship is provided by the Graduate School, part of the Office of Vice Chancellor for Research and Graduate Education at the University of Wisconsin-Madison, with funding from the Wisconsin Alumni Research Foundation and UW-Madison. L.F. was supported in part by the National Institute of General Medical Sciences of the National Institutes of Health under Award Number T32GM008505 (Chemistry–Biology Interface Training Program). L.F. was also supported by the 2024 Eli Lilly and Company/ACS Analytical Graduate Fellowship. L.F. acknowledges a predoctoral fellowship supported by the National Institutes of Health, under Ruth L. Kirschstein National Research Service Award from the National Institutes of Health-General Medical Sciences (F31GM156104). L.L. would like to acknowledge NIH grants RF1AG052324, R01AG078794, R21AG065728, S10OD028473, and S10OD025084, as well as funding support from a Vilas Distinguished Achievement Professorship and Charles Melbourne Johnson Professorship with funding provided by the Wisconsin Alumni Research Foundation and University of Wisconsin-Madison School of Pharmacy. The content is solely the responsibility of the authors and does not necessarily represent the official views of the National Institutes of Health or National Science Foundation. Figures 1 and Supplemental Figures 1-3 were generated in BioRender.

## Supplementary Data

Supplementary data are available at *Bioinformatics* online.

