## Supplemental File 1 for "EndoGenius: Enabling comprehensive identification and quantitation of endogenous peptides"

### Table of Contents

Supplemental Figures (located within this document)


- **Figure S1:** Integration of MotifQuest with EndoGenius
- **Figure S2:** Utilization of EndoGenius for quantitation of label-free and isobarically labeled samples
- **Figure S3:** Workflow for construction of spectral libraries with EndoGenius
- **Figure S4:** EndoGenius main GUI
- **Figure S5:** Available post-translational modifications for EndoGenius
- **Figure S6:** Supplemental tools for EndoGenius

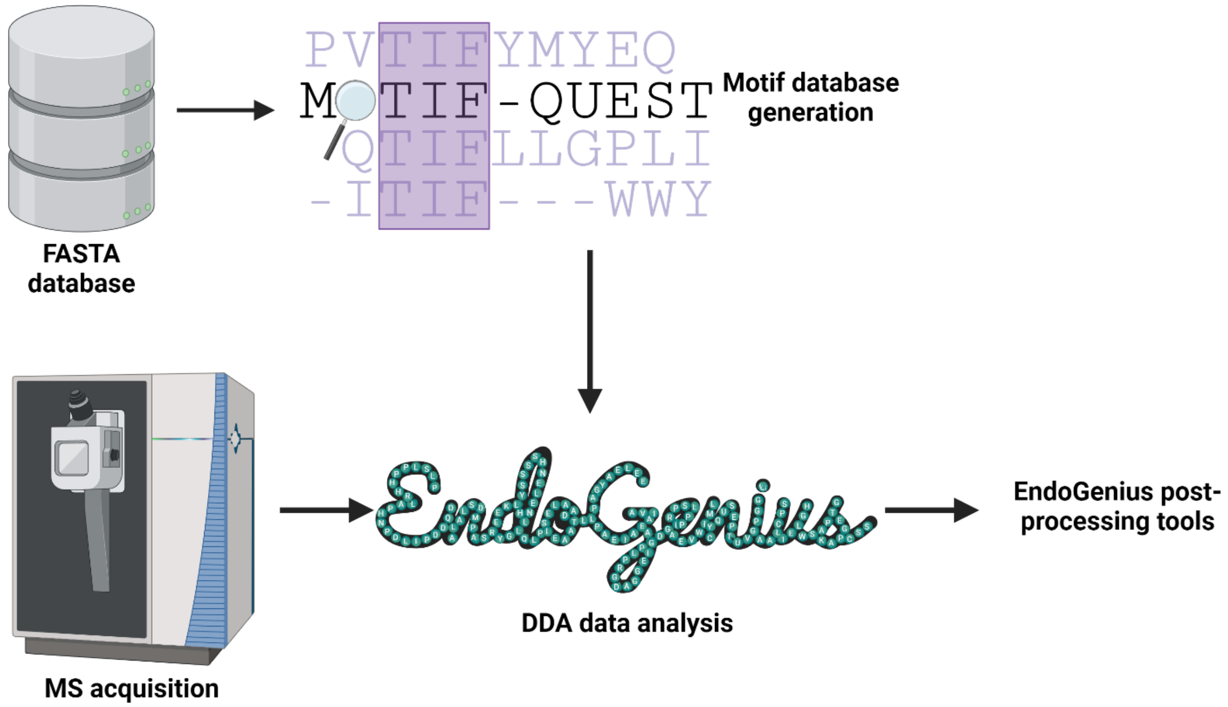


**Supplemental Figure 1:** Integration of MotifQuest software with the EndoGenius algorithm. A standard FASTA database file is provided as input to the MotifQuest algorithm.(Dang, et al., 2024) The output motif database is used within the motif evaluation stage of the EndoGenius algorithm. Following use of EndoGenius, incorporating post-processing tools, including quantification, can be applied.

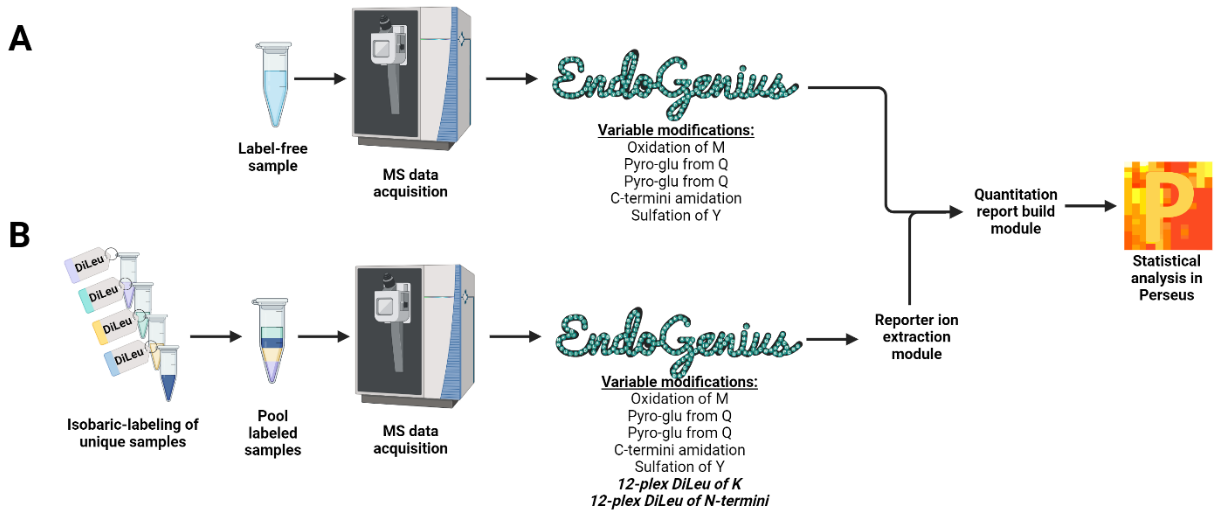


**Supplemental Figure 2:** Quantification with EndoGenius. **(A)** For label-free samples, following MS data acquisition, EndoGenius can be operated with a variety of post-translational modifications (PTMs). Following EndoGenius, a quantitation report can be built using the specified module, which generates a report compatible with Perseus software.(Tyanova, et al., 2016-06-

27) **(B)** For isobarically-labeled samples, the separately labeled samples are pooled and undergo MS data acquisition. When moving to EndoGenius, 12-plex DiLeu is selected as a variable modification on lysine and on the peptide N-termini. A specific reporter ion extraction module is included, following which the exported results can be imported into Perseus for further statistical analysis.

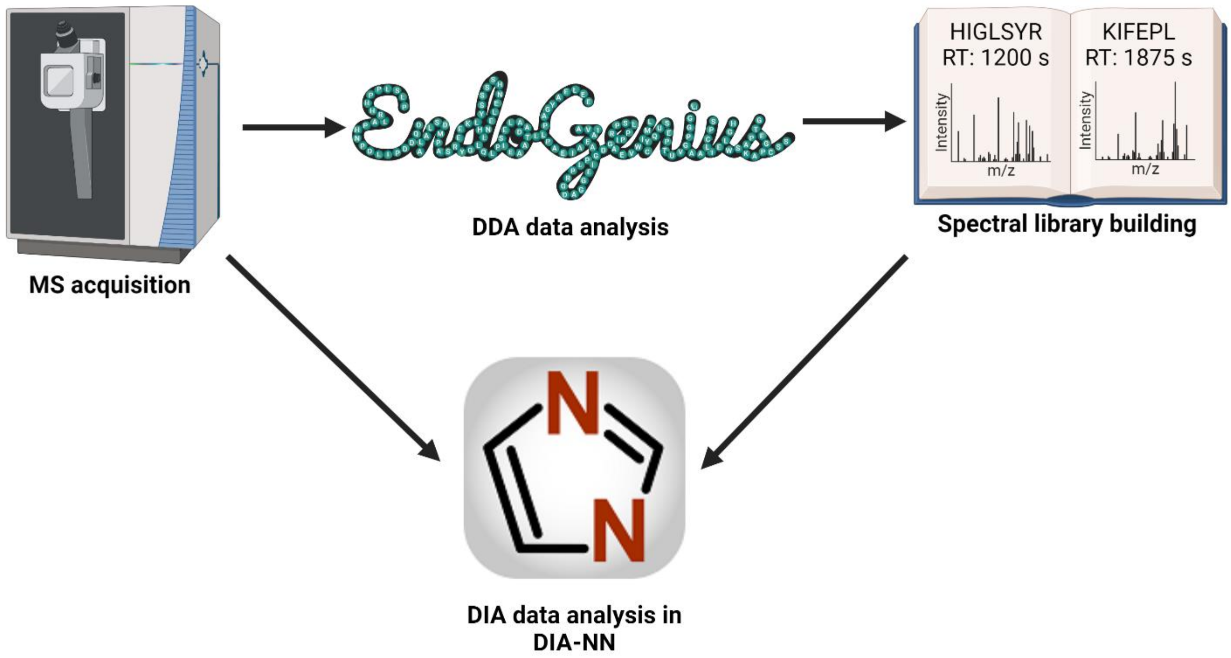


**Supplemental Figure 3:** The exported results from EndoGenius can be utilized to build a high-confidence spectral library. This spectral library is formatted to be compatible with DIA-NN software. DIA-NN can then be used to reference the spectral library for analysis of data-independent acquisition (DIA) spectral datasets.(Demichev, et al., 2020)

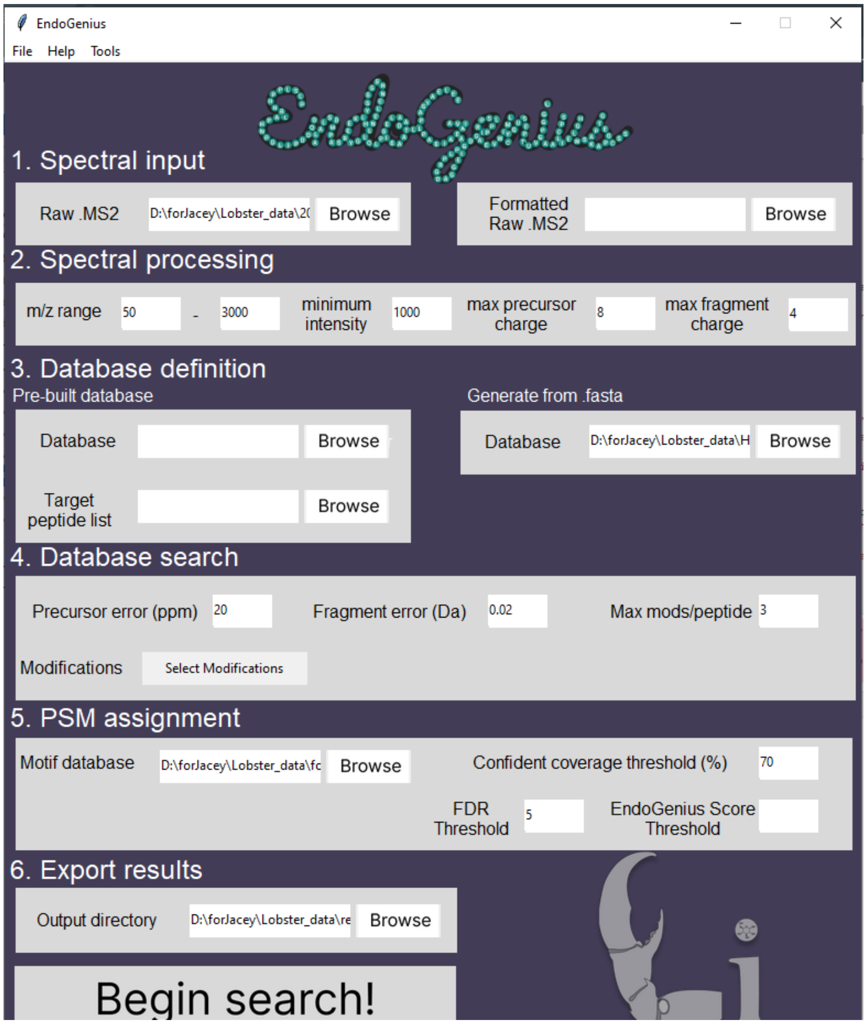


**Supplemental Figure 4:** Homepage graphical user interface (GUI) of EndoGenius. This includes all the necessary parameters to conduct an analysis within EndoGenius.

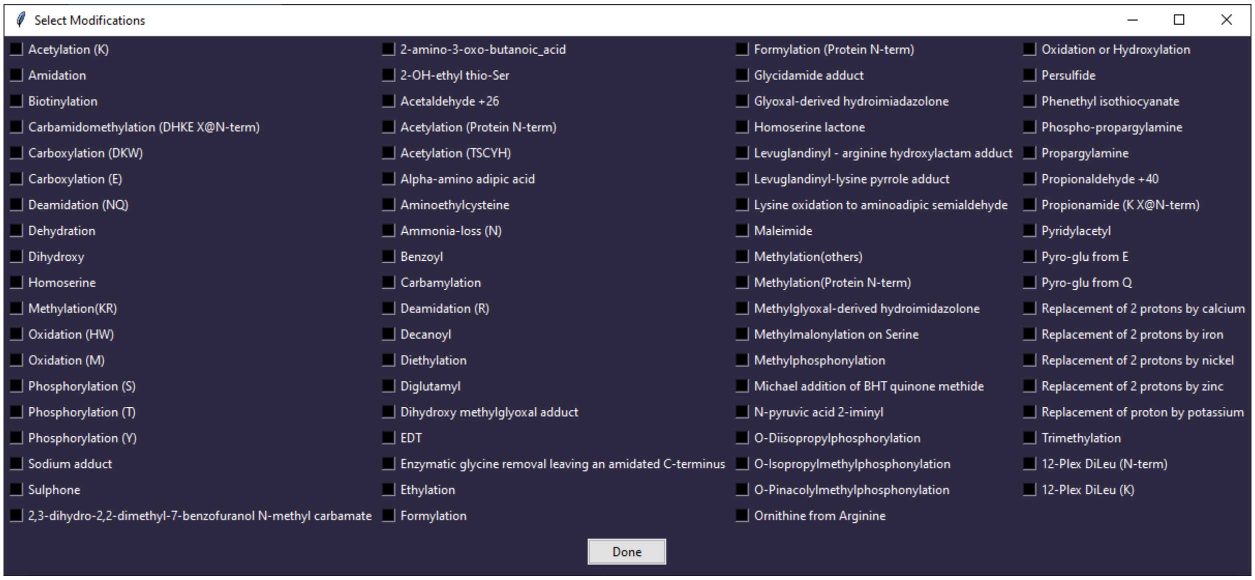


**Supplemental Figure 5:** Extended list of modifications available for selection in EndoGenius.

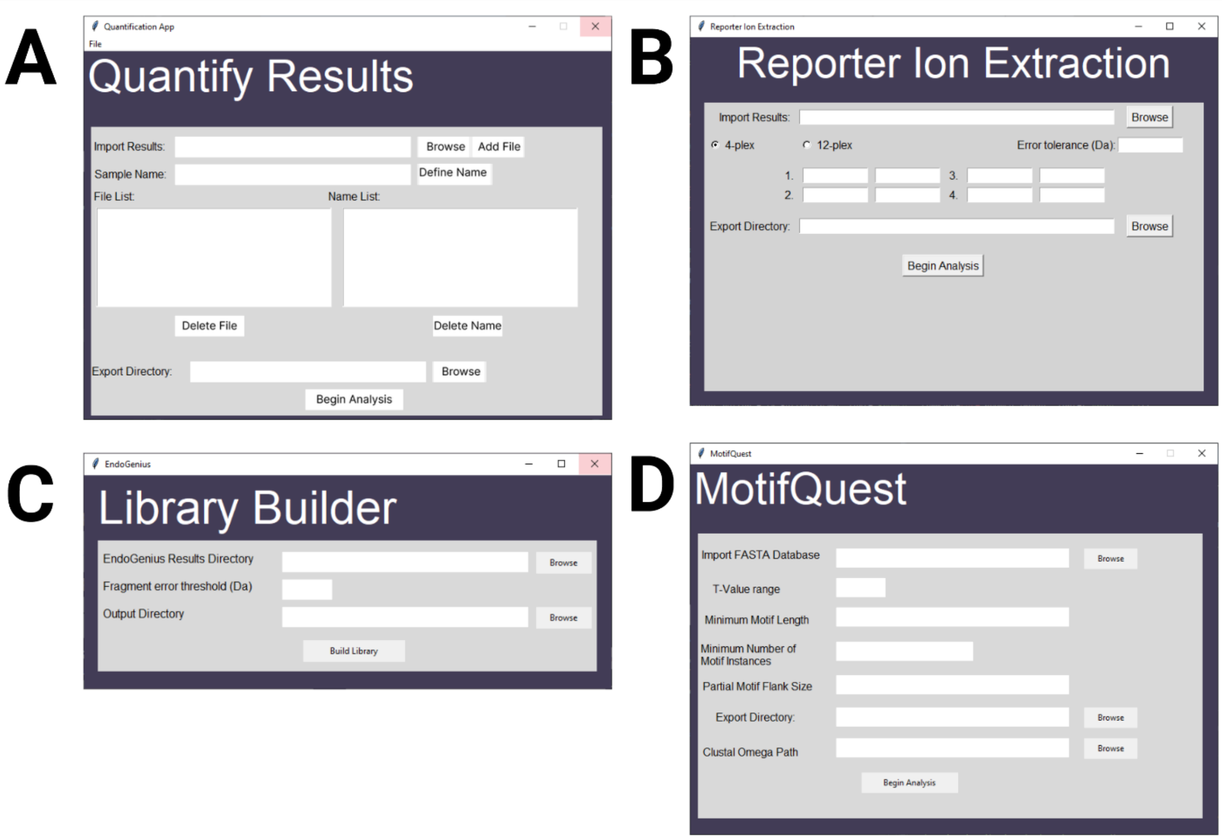


**Supplemental Figure 6:** Respective GUIs for tools integrated into EndoGenius. **(A)** Quantification of EndoGenius output across any number of samples. **(B)** Reporter ion extraction for 4-plex or 12-plex isobarically-labeled samples following EndoGenius search. **(C)** Library building for utility in searching DIA spectra. **(D)** MotifQuest integration for building of a custom motif database from a peptide sequence database.
